# Arabidopsis ENHANCED DISEASE RESISTANCE1 Protein Kinase Regulates the Association of ENHANCED DISEASE SUSCEPTIBILITY1 and PHYTOALEXIN DEFICIENT4 to Inhibit Cell Death

**DOI:** 10.1101/806802

**Authors:** Matthew Neubauer, Irene Serrano, Natalie Rodibaugh, Deepak D. Bhandari, Jaqueline Bautor, Jane E. Parker, Roger W. Innes

## Abstract

ENHANCED DISEASE SUSCEPTIBILITY1 (EDS1) and PHYTOALEXIN DEFICIENT4 (PAD4) are sequence-related lipase-like proteins that function as a complex to regulate defense responses in Arabidopsis by both salicylic acid-dependent and independent pathways. Here we describe a gain-of-function mutation in PAD4 (S135F) that enhances resistance and cell death in response to infection by the powdery mildew pathogen *Golovinomyces cichoracearum.* The mutant PAD4 protein accumulates to wild-type levels in Arabidopsis cells, thus these phenotypes are unlikely to be due to PAD4 over accumulation. The phenotypes are similar to loss of function mutations in the protein kinase Enhanced Disease Resistance1 (EDR1), and previous work has shown that loss of *PAD4* or *EDS1* suppresses *edr1*-mediated phenotypes, placing these proteins downstream of *EDR1*. Here we show that EDR1 directly associates with EDS1 and PAD4 and inhibits their interaction in yeast and plant cells. We propose a model whereby EDR1 negatively regulates defense responses by interfering with the heteromeric association of EDS1 and PAD4. Our data indicate that the S135F mutation likely alters an EDS1-independent function of PAD4, potentially shedding light on a yet unknown PAD4 signaling function.

## INTRODUCTION

Loss-of-function mutations in the *ENHANCED DISEASE RESISTANCE1* (*EDR1*) gene of Arabidopsis confer enhanced resistance to the powdery mildew pathogen *Golovinomyces cichoracearum* (Frye and Innes 1998). This enhanced resistance is correlated with enhanced cell death at the site of infection. The *edr1-1* mutation causes a premature stop codon in the *EDR1* gene, which encodes a protein kinase with homology to mitogen–activated protein kinase kinase kinases (MAPKKKs) belonging to the Raf family (Frye et al. 2001). The *edr1* mutant does not display constitutive expression of defense genes in the absence of a pathogen, indicating that the enhanced resistance is not caused by constitutive activation of systemic acquired resistance (Frye and Innes 1998); however, *edr1*-mediated disease resistance is suppressed by mutations that block or reduce salicylic acid (SA) production or signaling (Frye and Innes 1998; Frye et al. 2001; Christiansen et al. 2011; Hiruma et al. 2011; Hiruma and Takano 2014; Tang 2005), suggesting that *edr1*-mediated enhanced resistance against *G. cichoracearum* requires an intact SA signaling pathway.

In addition to enhancing resistance to powdery mildew, loss-of-function mutations in *EDR1* enhance drought-induced growth inhibition, ethylene induced senescence and sensitivity to abscisic acid (ABA) (Tang et al. 2005; Wawrzynska et al. 2008). The enhanced drought-induced growth inhibition and enhanced ABA sensitivity phenotypes, but not ethylene-induced senescence, are suppressed by mutations in the *ENHANCED DISEASE SUSCEPTIBILITY 1* (*EDS1*) and *PHYTOALEXIN DEFICIENT 4* (*PAD4*) genes, which encode sequence-related nucleocytoplasmic lipase-like proteins (Tang 2005). The inability of these mutations to suppress the ethylene-induced senescence phenotype of *edr1* mutants suggests that EDR1may regulate multiple pathways.

The *pad4* mutant was originally isolated in an Arabidopsis screen for enhanced disease susceptibility to *Pseudomonas syringae* pv. *maculicola* (Glazebrook and Ausubel 1994). PAD4 physically interacts with EDS1 as a heterodimer (Feys et al. 2001; Jirage et al. 1999; Rietz et al. 2011; Wagner et al. 2013), forming a nucleo-cytoplasmic complex that promotes accumulation of the plant defense signaling molecule SA (Cui et al. 2017; Feys et al. 2001; 2005). EDS1 and PAD4 also contribute to defense responses activated by intracellular nucleotide-binding, leucine rich repeat (NLR) receptors that have an N-terminal Toll-interleukin 1 receptor (TIR) domain (Aarts et al. 1998; Bhandari et al. 2019; Cui et al. 2018; Feys et al. 2001; Glazebrook and Ausubel 1994). NLR-mediated immune responses are often associated with localized host-cell death as part of the hypersensitive response (HR) (Maekawa et al. 2011). Arabidopsis *pad4* mutants display a delayed HR against the oomycete pathogen *Hyaloperospora arabidopsidis* that is insufficient for preventing pathogen spread (Feys et al. 2001). This partially retained HR can be attributed to partial genetic redundancy between *PAD4* and the nuclear *SENESCENCE-ASSOCIATED GENE 101* (*SAG101*), another component of the EDS1 regulatory hub (Feys et al. 2005; Lipka et al. 2005). It was recently established that EDS1-SAG101 heterodimers promote HR cell death in TIR-NLR receptor immunity, whereas formation of EDS1-PAD4 heterodimers is necessary for transcriptionally mobilizing SA and other defense pathways (Bhandari et al. 2019; Feys et al. 2005; Gantner et al. 2019; Lapin et al., 2019; Rietz et al. 2011). Complementary studies have shown that EDS1 and PAD4 transduce photo-oxidative stress signals leading to cell death and the slowing of plant growth, and that they are involved in plant fitness regulation (Chandra-Shekara et al. 2007; Venugopal et al. 2009; Wituszynska et al. 2013; Xiao et al. 2001).

So far, all described mutations in *EDS1* and *PAD4* have caused a loss of function (Feys et al. 2001; Glazebrook 1999; Hu et al. 2005; Jirage et al. 1999; Rietz et al. 2011; Wagner et al. 2011). Here we describe a gain-of-function mutation in the *PAD4* gene that enhances a subset of *edr1* mutant phenotypes, including *edr1*-dependent cell death after powdery mildew infection, and *edr1* accelerated ethylene- and age-induced senescence. This mutation causes a serine to phenylalanine substitution at position 135 of PAD4. Furthermore, the PAD4^S135F^ substitution alone confers enhanced disease resistance and enhanced cell death after infection with the powdery mildew fungus *G. cichoracearum*. The molecular basis for these phenotypes remains unclear, however, the S135F substitution did not affect PAD4 protein accumulation, localization, or its ability to associate with EDS1. The discovery that *pad4*^*S135F*^ enhances a subset of *edr1* phenotypes supports previous findings that the *edr1* phenotype is at least partially due to changes in SA signaling (Tang et al. 2005). Analysis of *edr1* and *pad4*/*eds1* transcriptome data revealed that a significant proportion of the PAD4/EDS1 gene network is upregulated in *edr1* plants during the defense response. To follow up on these results, we investigated whether EDR1 plays a direct role in regulating PAD4. Significantly, we found that EDR1 interacts with both PAD4 and EDS1, and that EDR1 can inhibit the interaction between EDS1 and PAD4.

## RESULTS

### Identification of a mutation in *PAD4* that enhances *edr1* mutant phenotypes

The *edr1* mutant displays enhanced sensitivity to *flg22*, a 22 amino-acid peptide derived from bacterial flagellin that is known to induce defense responses (Geissler et al. 2015). This sensitivity can be assayed in very young seedlings grown in liquid culture. We took advantage of this phenotype to screen for second site mutations that can suppress this enhanced flg22 sensitivity, restoring *edr1* mutants to a wild-type phenotype. Candidate suppressor mutants obtained in this screen were assessed for the presence of mutations in genes previously shown to be required for *edr1* mutant phenotypes (Tang 2005; Wawrzynska et al. 2008), so that we could focus our efforts on new genes. To our surprise, all suppressor candidates analyzed (13 in total) carried an identical missense mutation in the *PAD4* gene, causing a change of amino acid Ser135 to Phe135 (*pad4*^*S135F*^). Because these 13 mutants were derived from multiple different EMS-mutagenized parents, it seemed likely that the parent population (prior to mutagenesis) carried this mutation, and that the mutation was not responsible for the suppressor phenotype. We therefore sequenced the *PAD4* gene in the *edr1-1* parental line used for suppressor mutagenesis. This analysis confirmed that the *edr1-1* parental line used for the suppressor mutagenesis carried the same mutation, and that this mutation had arisen at some point during the backcrossing process of the original *edr1-1* mutant, which lacks this mutation (see Methods).

### The *pad4*^*S135F*^ mutation confers enhanced disease resistance and contributes to *edr1*-dependent enhanced cell death

Because we had previously shown that loss-of-function mutations in *PAD4* suppressed *edr1-1* mutant phenotypes (Tang 2005), the discovery that a missense mutation in *PAD4* was present in the *edr1-1* mutant suggested that the *pad4*^*S135F*^ mutation might be contributing to *edr1* mutant phenotypes. To test this hypothesis, we infected wild-type Col-0, *edr1-1*, *edr1*-*3* (contains a T-DNA insertion in *EDR1*), *pad4*^*S135F*^, and *edr1-1 pad4*^*S135F*^ plants with *G. cichoracearum* and quantified fungal growth by counting conidiospores at 8 dpi. As expected, *edr1-1 pad4*^*S135F*^ plants had a reduced spore count compared to wild-type Col-0 (Fig. 1A). This enhanced disease resistance was not influenced by the presence of the *pad4*^*S135F*^ mutation, as the *edr1-1* and *edr1-3* mutants had comparable spore counts (Fig. 1A). Interestingly, the *pad4*^*S135F*^ mutant also had a reduced spore count, similar to that of the *edr1* mutants (Fig. 1A). These results indicate that the *pad4*^*S135F*^ mutation alone confers an enhanced disease resistance similar to *edr1* mutations, and that the mutations are not additive in their effects.

**Fig. 1.**
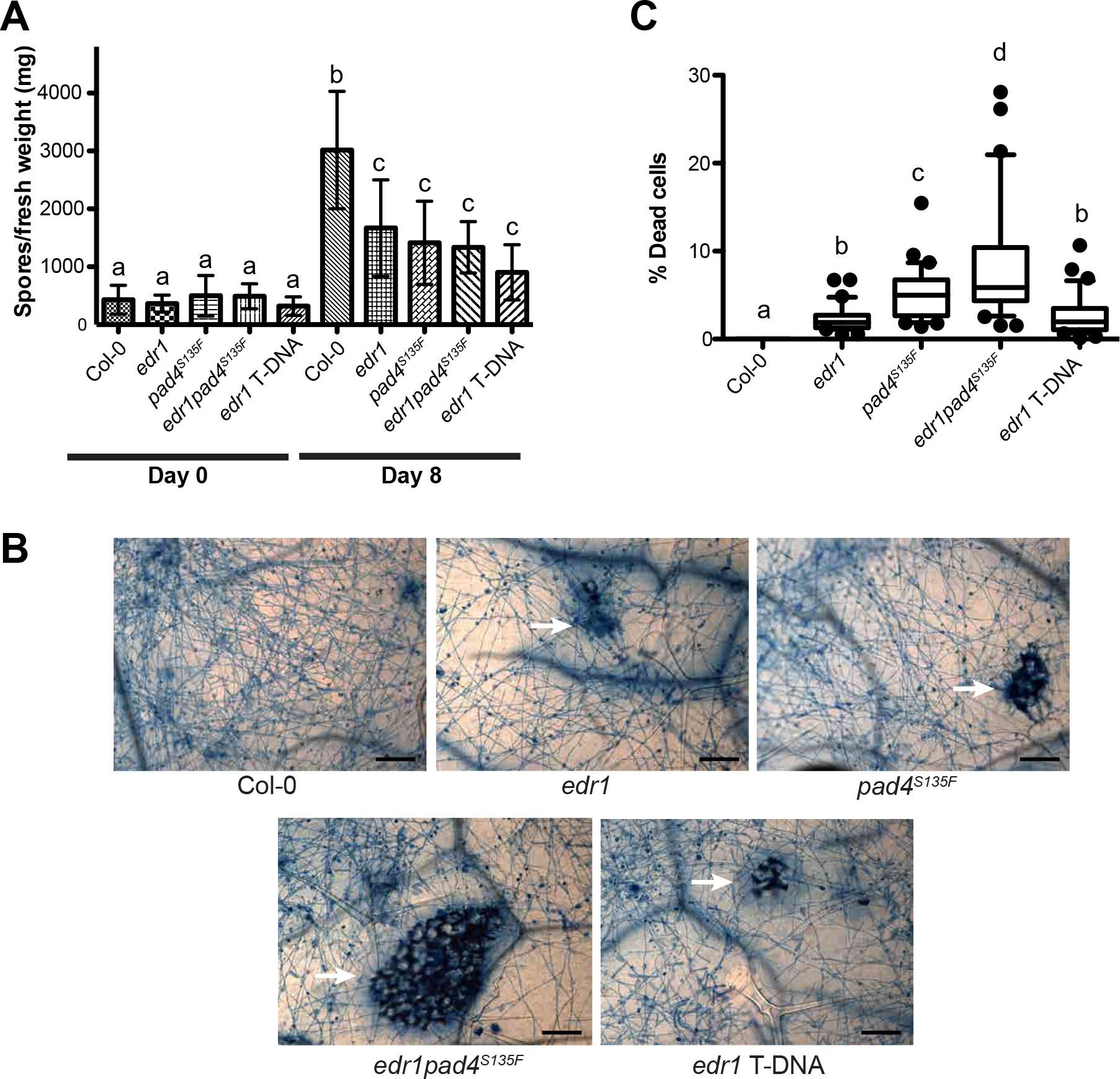
The *pad4*^*S135F*^ mutation confers enhanced disease resistance and contributes to *edr1*-associated cell death. **A**, Quantitative analysis of powdery mildew conidia (asexual spores) on Col-0, *edr1-1*, *pad4*^*S135F*^, *edr1-1pad4*^*S135F*^ and *edr1-3* lines. Plants were inoculated with powdery mildew and conidia production was determined 8 dpi. Bars indicate the mean of three samples, each with three technical replicates. Error bars indicate SD. Results are representative of 3 independent experiments. **B**, trypan blue staining of powdery mildew-infected Col-0, *edr1-1*, *pad4*^*S135F*^, *edr1-1pad4*^*S135F*^ and *edr1-3* lines. The indicated lines were assessed for leaf mesophyll cell death 8 dpi and cell death was quantified using ImageJ. For quantification, six pictures from five independent experiments were randomly chosen (n=30). Results are provided as means with 10th and 90th percentiles (box) and range (whiskers). Statistical outliers are shown as a circle. Lower case letters indicate values that are significantly different (P<0.01; one-way ANOVA test using the Bonferroni method). **C**, Four-week old plants were infected with *G. cichoracearum* and phenotypes were scored 8 days post-infection. Trypan blue staining of infected leaves to reveal fungal hyphae and patches of dead mesophyll cells (arrows). Bars=50 μm. Pictures are representative of 3 independent experiments

Loss-of-function mutations in *PAD4* have been shown to enhance disease susceptibility (Feys et al. 2001; Frye et al. 2001; Glazebrook et al. 1997; Zhou et al. 1998). Indeed, upon *G. cichoracearum* infection, *pad4-1* plants accumulate more fungal spores than wild-type (Supplementary Fig. S1). These data indicate that the *pad4*^*S135F*^ mutation causes a gain-of-function that enhances resistance to *G. cichoracearum*.

In addition to enhancing resistance to *G. cichoracearum*, the *edr1* mutation causes an increase in mesophyll cell death following infection by this fungus (Frye and Innes 1998). To assess whether the *pad4*^*S135F*^ mutation contributes to this cell death phenotype, we used trypan blue staining to score cell death at 5 dpi. The *edr1-1 pad4*^*S135F*^ mutant displayed large patches of mesophyll cell death (Fig. 1B). In comparison, the *edr1-1* and *edr1-3* mutants displayed fewer patches of dead cells, and these patches were smaller. Significantly, the *pad4*^*S135F*^ mutant also displayed patches of dead mesophyll cells, similar in appearance to the *edr1* mutants. No mesophyll cell death was detected in wild-type Col-0 plants. To further characterize the cell death response, the patches of dead mesophyll cells positive for trypan blue staining were quantified. The *edr1*-dependent cell death was enhanced by the presence of the *pad4*^*S135F*^ mutation, indicating that the two mutations are additive in their effect on powdery mildew-induced cell death (Fig. 1C). Notably, *pad4*^*S135F*^ plants displayed a significantly higher level of cell death than *edr1* plants.

### EDR1 physically interacts with EDS1 and PAD4

The conclusion that *pad4*^*S135F*^ can enhance some but not all *edr1* phenotypes prompted us to investigate whether EDR1 and PAD4 are part of a common regulatory complex. In support of this hypothesis, both proteins were previously shown to localize partially to the nucleus (Feys et al. 2005; Christiansen et al. 2011). To test whether EDR1 interacts with PAD4, we performed yeast two-hybrid analyses. Counter to expectations, we could not detect an interaction between wild-type EDR1 and PAD4 (Fig. 2A). As described above, however, PAD4 is known to interact with EDS1, and this interaction is required for both basal disease resistance and TIR-NLR-mediated resistance (Feys et al. 2005; 2001; Rietz et al. 2011; Wagner et al. 2013), suggesting that the genetic interaction between *EDR1* and *PAD4* could be mediated by EDS1. We thus tested whether EDR1 interacts with EDS1, and observed a positive yeast two-hybrid interaction (Fig. 2A). One possible reason we could not detect the interaction between PAD4 and EDR1 is that PAD4 could be a substrate of EDR1, and this interaction may be very transient. We therefore tested whether a substrate-trap mutant form of EDR1, EDR1^D810A^ (Gu and Innes 2011), interacts with PAD4. Indeed, EDR1^D810A^ was found to interact with both EDS1 and PAD4. However, the enhanced interaction of EDR1^D810A^ with PAD4 is possibly explained by enhanced stability of the mutant protein compared to wild-type EDR1 (Fig. 2A).

**Fig. 2.**
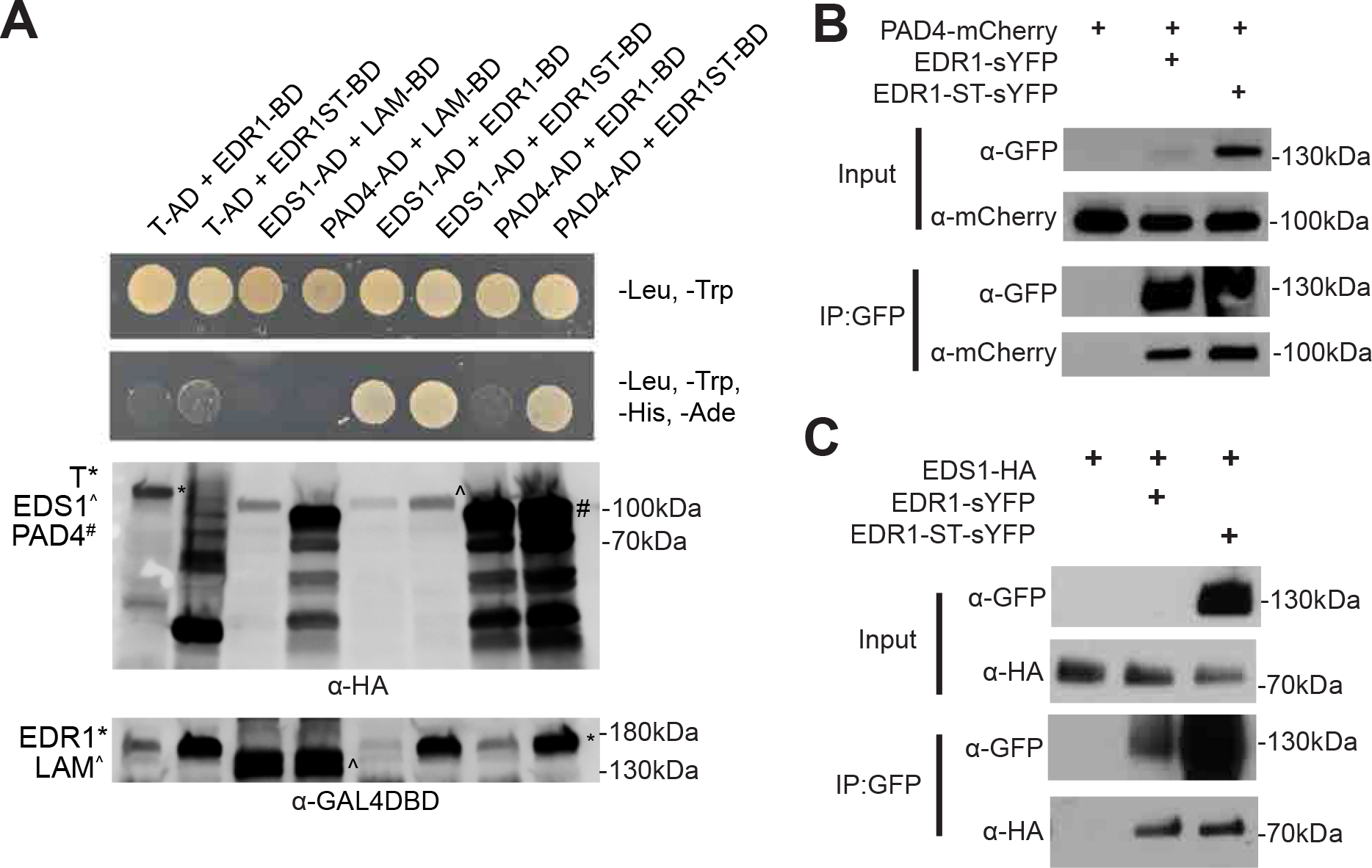
EDR1 physically interacts with EDS1 and PAD4. **A,** Yeast two-hybrid analysis of EDR1 interactions with EDS1 and PAD4. AD, GAL4 activation domain fusion; BD, GAL4 DNA binding domain fusion; T, SV40 large T antigen; LAM, lamin. Protein expression was verified through immunoblotting. AD-tagged proteins also contain an HA tag, which was used for detection. **B,** EDR1 co-immunoprecipitates with PAD4. **C,** EDR1 co-immunoprecipitates with EDS1. For both panels B and C, the indicated constructs were transiently expressed in *N. benthamiana* and then immunoprecipitated using GFP-Trap beads. Note that wild-type EDR1-sYFP accumulates poorly when transiently expressed in *N. benthamiana*, but can still be immunoprecipitated in sufficient levels. These experiments were all repeated three times with similar results.

We then sought to determine whether the interactions observed in yeast also occur *in planta*. Co-immunoprecipitation (Co-IP) assays in *N. benthamiana* were performed. EDS1-3xHA and PAD4-mCherry were independently co-expressed with either an empty vector negative control, EDR1-sYFP, or EDR1^ST^-sYFP. Both PAD4 and EDS1 were found to Co-IP with EDR1 and EDR1^ST^, but not when co-expressed with an empty vector (Fig. 2B, 2C). These assays indicate that both PAD4 and EDS1 can form complexes with EDR1 and EDR1^ST^ *in planta.* As we observed in yeast, the EDR1^ST^ protein accumulated to higher levels than wild-type EDR1 (Fig. 2B and 2C). However, similar levels of EDS1 and PAD4 co-immunoprecipated with EDR1 and EDR1^ST^ (Fig. 2B and 2C). Based on these observations, we propose that EDR1 directly interacts with both EDS1 and PAD4.

### EDR1 Inhibits the Interaction between EDS1 and PAD4

The interaction between EDR1 and both PAD4 and EDS1 raised the question of whether EDR1 regulates PAD4-EDS1 heterodimer association. Formation of the EDS1-PAD4 heterodimer brings together α-helical coil surfaces in the partner C-terminal EP-domains that are essential for basal and TIR-NLR immunity signaling (Bhandari et al., 2019; Lapin et al., 2019). To test whether EDR1 can affect this interaction, we performed a yeast three-hybrid analysis in which the kinase domain of EDR1 (EDR1-KD) was expressed as a third protein in the yeast cell under control of the methionine-regulated promoter Met25 (repressed in the presence of 1 mM methionine and induced in its absence). However, we still observed accumulation of EDR1-KD in the absence of methionine, perhaps due to leakiness of the promoter (Fig. 3). EDR1-KD expression inhibited the interaction between EDS1 and PAD4 (Fig. 3A). To test whether this effect of EDR1 was dependent on EDR1 kinase activity, we also performed the assay using EDR1-KD^ST^, which is kinase-inactive. EDR1-KD^ST^ also blocked the EDS1-PAD4 interaction (Fig. 3A). Expression of EDR1-KD and EDR1-KD^ST^ had no noticeable effect on the interaction between the bacterial effector AvrB and the soybean R protein RIN4b, indicating that the effect on the EDS1-PAD4 interaction was specific. Immunoblotting demonstrated that EDR1-KD and EDR1-KD^ST^ accumulated in yeast to similar levels, and that EDR1 expression did not interfere with the accumulation of EDS1 or PAD4 (Fig. 3B). That EDR1 kinase activity was dispensable for blocking the EDS1-PAD4 interaction suggests that EDR1 may be interfering with EDS1-PAD4 association by competing for a common EDS1 binding site, rather than by phosphorylation of either protein.

**Fig. 3.**
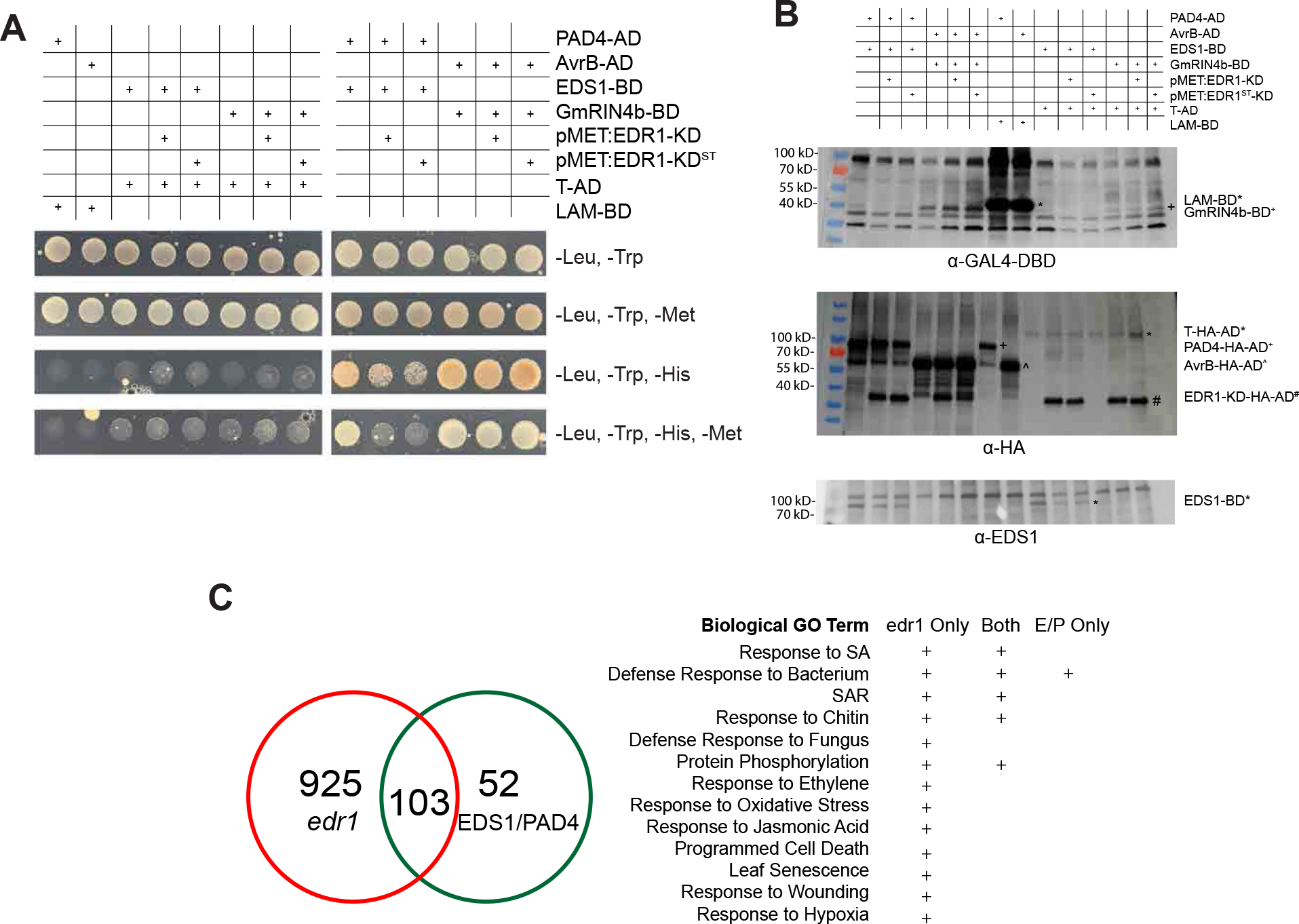
EDR1 interferes with EDS1:PAD4 association. **A,** The EDR1 kinase domain (KD) inhibits EDS1:PAD4 interaction in a yeast three-hybrid assay. The indicated constructs were transformed into yeast strains AH109 (activation domain constructs) and Y187 (DNA binding domain and methionine promoter constructs in pBridge vector) and then mated. Diploids were selected on minus Leu Trp plates, then replated on the indicated media. Growth on minus His plates indicates physical interaction between EDS1 and PAD4. Media lacking methionine induces the MET promoter. AvrB and RIN4b are positive controls for interaction. **B**, Immunoblot analysis confirms protein expression in yeast strains utilized in yeast three-hybrid assay. **C**, Loss of *EDR1* results in the upregulation of the EDS1-PAD4 network during a defense response. The *edr1* only dataset is enriched for a more diverse set of biological GO terms than the EDS1-PAD4 network.

### *edr1* Plants Display Enhanced EDS1/PAD4 Signaling During Defense Response

Recently, a network of 155 core genes was demonstrated to be upregulated during the overexpression of EDS1 with PAD4 (Cui at al. 2017). Previous work has demonstrated that loss of function mutations in either *EDS1* or *PAD4* inhibit a subset of *edr1* phenotypes (Tang 2005). The discovery that EDR1 can interact with EDS1 and PAD4, as well as disrupt the formation of the EDS1/PAD4 complex, prompted us to investigate whether EDR1 negatively regulates the EDS1-PAD4 signaling network. We have previously demonstrated that the loss of *EDR1* results in the upregulation of many defense-related genes during powdery mildew infection (Christiansen et al. 2011). We found that the majority of the 155 genes that were upregulated during EDS1-PAD4 overexpression are significantly upregulated in *edr1* plants relative to wildtype after powdery mildew infection (Fig. 3C). 103 of the 155 EDS1-PAD4 upregulated transcripts were upregulated in *edr1* plants during infection. This demonstrates that EDR1 has a negative impact on the induction of many EDS1-PAD4 upregulated genes during the defense response.

GO term enrichment analysis revealed that the genes belonging to both the EDS1-PAD4 upregulated and *edr1* upregulated networks are enriched for processes such as SA response, response to chitin, and protein phosphorylation (Fig. 3C). Interestingly, those genes that were found to be upregulated in *edr1* plants, but not belonging to the EDS1-PAD4 network, were enriched for a more diverse set of processes, including response to JA, ethylene, oxidative stress, hypoxia, and wounding. This correlates with the previous discovery that *edr1* phenotypes are only partially supressed by mutations in *EDS1* or *PAD4* (Tang 2005), as well as the observation that *pad4*^*S135F*^ enhances a subset of *edr1* phenotypes (Fig. 1). These data demonstrate that EDR1 negatively regulates a broad set of defense responses, which includes but is not limited to, the EDS1-PAD4 network.

### The *pad4*^*S135F*^ mutation does not affect protein accumulation, localization, or interaction with EDS1

To determine the effect of the *pad4*^*S135F*^ mutation on PAD4 function, we investigated possible changes that could result in PAD4 over-activity. We hypothesized that an increase in the stability of the PAD4 protein caused by the *pad4*^*S135F*^ mutation might result in enhanced SA signaling and cell death. However, we were unable to detect an increase in the accumulation of PAD4^S135F^ relative to PAD4 in Arabidopsis plants undergoing a defense response elicited by the RPS4 TIR-NLR protein (unelicited plants have nearly undetectable levels of PAD4; Fig. 4A).

**Fig. 4.**
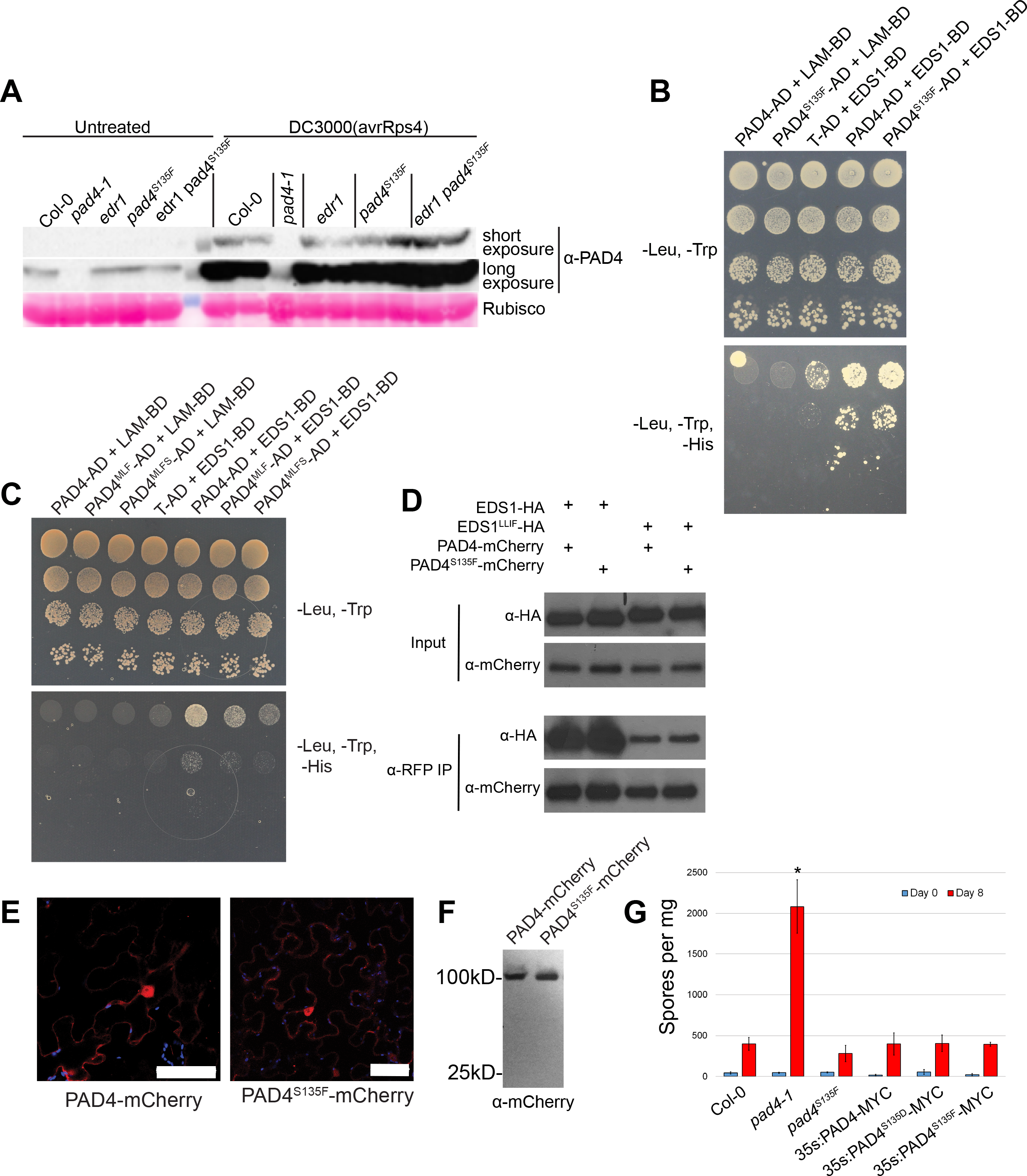
The S135F mutation in PAD4 does not affect its stability, interaction with EDS1, or subcellular localization pattern. **A,** PAD4 protein accumulates to similar levels in wild-type Col-0, pad4S135F, edr1 and double mutant Arabidopsis. Total protein was extracted from Arabidopsis rosette leaves that were either untreated or sprayed with Pseudomonas syringae DC3000(avrRps4), which induces PAD4 accumulation. **B,** PAD4S135F interacts with EDS1 in a yeast two-hybrid assay. The indicated constructs were transformed into yeast strain AH109 (activation domain constructs, AD) and yeast strain Y187 (DNA binding domain constructs, BD) and the strains mated, with diploids plated on the indicated media. **C,** The S135F mutation does not enhance the ability of PAD4MLF to interact with EDS1 in a yeast two-hybrid assay. The indicated constructs were transformed into yeast strain AH109 (activation domain constructs, AD) and yeast strain Y187 (DNA binding domain constructs, BD) and the strains mated, with diploids plated on the indicated media. **D,** The S135F mutation does not increase the interaction between PAD4 and EDS1LLIF. Constructs were expressed in N. benthamiana and protein immunoprecipitated using anti-RFP beads. **E,** PAD4S135F displays a nucleocytoplasmic localization pattern indistinguishable from wild-type PAD4. The indicated constructs were transiently expressed in N. benthamiana and imaged using confocal microscopy. Scale bar = 50 μM. **F,** PAD4-mCherry and PAD4S135F-mCherry accumulate at similar levels without free mCherry tag. Tissue from E was subjected to immunoblotting using an anti-mCherry antibody. **G,** PAD4^S135D^ and PAD4^S135F^ both can complement a *pad4-1* loss of function mutation. Four week old Arabidopsis plants were infected with powdery mildew. Spore counts were taken immediately following infection and 8 dpi. Bars indicate the means ±SD of three biological replicates per genotype. Asterisk denotes P value < 0.05 (Student’s T-test for pairwise comparisons to all other genotypes).

Another possible explanation for the over-activity of PAD4^S135F^ is that it might have enhanced interaction with its partner, EDS1. The EDS1-PAD4 interaction is mediated principally by conserved residues in the partner N-terminal domains, respectively EDS1^LLIF^ and PAD4^MLF^ that form a hydrophobic groove (Wagner et al. 2013). In an Arabidopsis EDS1-PAD4 structural model based on the EDS1-SAG101 heterodimer crystal structure (Wagner et al. 2013), PAD4^S135^ is located in a loop close to, but facing away from the PAD4^MLF^ heterodimer contact site (Supplementary Fig. S2). We therefore assessed whether the S135F substitution in PAD4 affected its interaction with EDS1 in a yeast two-hybrid assay. We observed no obvious effect on the interaction (Fig. 4B). In addition, we introduced the S135F mutation into the PAD4^MLF^ triple mutant, generating PAD4^MLFS^. We found that the S135F mutation did not significantly enhance the weakened interaction between PAD4^MLF^ and EDS1 in yeast two-hybrid assays (Fig. 4C). Similarly, we observed no change in the ability of PAD4^S135F^ to co-immunoprecipitate with EDS1 or with EDS1^LLIF^ compared to WT PAD4 (Fig. 4D). These data indicate that the S135F mutation does not affect the ability of PAD4 to interact with EDS1.

Finally, we investigated whether the S135F mutation alters the localization of PAD4 in plant cells. Transient expression of PAD4-mCherry and PAD4^S135F^-mCherry showed that both proteins displayed a nucleocytoplasmic localization (Fig. 4E). To verify that the observed localization was not the result of protein degradation, we performed immunoblotting, which also demonstrated a similar level of accumulation of the PAD4 and PAD4^S135F^ proteins (Fig. 4F). We thus conclude that the S135F mutation does not alter PAD4 stability, localization, or its ability to interact with EDS1, but somehow still affects PAD4 function and signaling.

### Phosphorylation of PAD4^S135^ is Unlikely to Negatively Regulate PAD4 Activity

Our data indicate that EDR1 functions as a negative regulator of EDS1/PAD4 signaling. As EDR1 has been demonstrated to have kinase activity (Tang and Innes 2002), we hypothesized that EDR1-mediated regulation of EDS1/PAD4 is by direct phosphorylation. Therefore, we carried out IP-MS experiments in *N. benthamiana* using transient expression of Arabidopsis PAD4, EDS1, EDR1, and EDR1^ST^ proteins. However, we were consistently unable to detect any phosphorylation of PAD4 or EDS1 in either the presence or absence of active EDR1. This result was repeated in three independent experiments. Importantly, the unphosphorylated S135-containing peptide was identified in all replicates, even though PAD4^S135^ is surface exposed in the structural model (Supplementary Fig. S2), making it potentially amenable for phosphorylation.

Although we could not detect EDR1-mediated phosphorylation of EDS1 or PAD4 in *N. benthamiana*, it remains a possibility that under specific conditions, EDR1 or some other kinase may regulate PAD4 via phosphorylation. Thus, we investigated whether the gain of function phenotype of S135F may be caused by the loss of an important phosphorylated serine residue. To test whether S135 is an important site of phosphorylation, we generated transgenic *pad4-1* PAD4^S135D^-MYC phosphomimic Arabidopsis. If PAD4 is indeed negatively regulated by phosphorylation at S135, then the PAD4^S135D^-MYC transgene should be unable to complement the *pad4-1* allele. However, we found that *pad4-1* plants were fully complemented by PAD4^S135D^-MYC, PAD4-MYC, and PAD4^S135F^-MYC expression in resistance to powdery mildew infection (Fig. 4G). This result demonstrates that the gain of function phenotype of S135F is unlikely to be the result of blocking phosphorylation.

## Discussion

Arabidopsis EDR1 acts as a negative regulator of cell death during both biotic and abiotic stress responses. Loss-of-function mutations in the *EDR1* gene confer enhanced disease resistance to powdery mildew infection and more rapid senescence than wild-type plants when exposed to ethylene (Frye and Innes 1998; Frye et al. 2001; Tang 2005). In this work, we report that a mutation in the *PAD4* gene (*pad4*^*S135F*^) enhances *edr1*-dependent cell death after pathogen attack. Moreover, the *pad4*^*S135F*^ mutation alone confers enhanced disease resistance to the powdery mildew *G. cichoracearum* and accelerated cell death.

PAD4 is required for the accumulation of the signaling molecule SA (Jirage et al. 1999; Feys et al. 2005), and thus loss-of-function mutations in the *PAD4* gene severely compromise defense against biotrophic pathogens, including powdery mildew (Glazebrook and Ausubel 1994; Gao et al. 2014). The *pad4*^*S135F*^ mutation, in contrast, enhances resistance to *G. cichoracearum*, indicating that this mutation causes a gain-of-function. Moreover, this enhanced disease resistance is accompanied by enhanced cell death (Fig. 1B), similar to that observed in the *edr1* mutant (Frye and Innes 1998). While the enhanced disease resistance is not additive in the *edr1-1pad4*^*S135F*^ double mutant, the cell death is more extensive in the double mutant than in either of the single mutants, suggesting that *PAD4* and *EDR1* independently regulate the cell death pathway.

The enhanced disease resistance phenotype in both *edr1* and *pad4*^*S135F*^ without additive effects in the double mutant can be explained by both mutations causing a similar effect on SA signaling. Alternatively, PAD4^S135F^ might be augmenting *edr1* cell death in parallel to SA, since PAD4 with EDS1 promotes both SA-dependent and SA-independent pathways in basal and TIR-NLR-mediated resistance (Cui, 2018; Bhandari 2019). We have shown that *pad4*^*S135F*^ does not alter PAD4 accumulation, localization, or interaction with EDS1 (Fig. 4), yet it remains unclear what effect this mutation has on PAD4. While PAD4^S135^ is located close to the chief N-terminal PAD4^MLF^ interface with EDS1^LLIF^, it is facing away from the interaction groove (Supplementary Fig. S2), consistent with the finding that the PAD4^S135F^ mutation does not obviously alter PAD4-EDS1 heterodimerization. It is possible that close proximity of PAD4^S135F^ to an α-helix of the PAD4 EP-domain (Supplemenatary Fig. S2) creates a loosening of N-terminal restraint on the PAD4 C-terminal signaling function. Recently, it has been demonstrated that EDS1/PAD4 functions to antagonize the activity of MYC2, a master regulator of JA signaling in TIR-NLR immunity (Cui et al., 2018). It is therefore a formal possibility that the S135F mutation alters the interaction between PAD4 and MYC2, or some other unknown signaling partner.

Although we could not detect an enhanced interaction between PAD4^S135F^ and EDS1 using a yeast two-hybrid assay, we did observe that co-expression of EDR1 with EDS1 and PAD4 inhibited the EDS1-PAD4 interaction in a yeast three-hybrid assay. Furthermore, EDR1 interacts strongly with EDS1 and PAD4 in yeast, and in co-IPs from *N. benthamiana*. Collectively, these observations suggest that EDR1 functions, at least in part, to negatively regulate the interaction between EDS1 and PAD4. Because formation of an EDS1-PAD4 heterodimer is essential for the rapid transcriptional reprogramming of host defense pathways in pathogen resistance (Bhandari et al. 2019), EDR1 might exert important negative control on EDS1-PAD4 signaling activity in response to infection. In support of this model, mutations in either *EDS1* or *PAD4* block *edr1*-mediated enhanced resistance and cell death (Frye et al. 2001). Furthermore, genes upregulated in the absence of EDR1 overlap significantly with genes upregulated by co-overexpression of EDS1 and PAD4 (Fig. 3C). Importantly, overexpression of either EDS1 or PAD4 alone does not upregulate these genes or enhance resistance (Cui et al. 2017), which indicates that it is the concentration of the EDS1- PAD4 complex, and not their individual protein levels, that determines the strength of defense signaling.

## MATERIALS AND METHODS

### Plant material and growth conditions

*Arabidopsis thaliana* accession Col-0, and Col-0 mutants *edr1-1* (Frye and Innes 1998), *edr1-3* (salk_127158C), *pad4*^*S135F*^, and *edr1-1 pad4*^*S135F*^ were used in this study. The *edr1-1* parental seed used for the suppressor mutagenesis was derived from a backcross 3 population. To confirm that the *pad4*^*S135F*^ mutation was present in this population, we sequenced *PAD4* amplified from multiple individuals of that population and found that the *pad4*^*S135F*^ mutation was segregating within the population. To assess whether the *pad4* ^*S135F*^ mutation was present in our original *edr1-1* mutant, we sequenced *PAD4* in an *edr1-1* M6 population (8 individual plants) that had never been backcrossed. Surprisingly, none of these plants carried the *pad4*^*S135F*^ mutation, suggesting that the mutation had arisen spontaneously at some point during the backcrossing process. Consistent with this conclusion, an *edr1-1* population being used by a former lab member in China also lacks this mutation (D. Tang, personal communication).

Seeds were surface sterilized with 50% (v/v) bleach and planted on one-half-strength Murashige and Skoog plates supplemented with 0.8% agar and 1% sucrose. Plates were placed at 4°C for 72 h for stratification and then transferred to a growth room set to 23°C and 9 h light (150 μEm-2s-1)/15 h dark cycle. Seven-day-old seedlings were transplanted to MetroMix 360 (Sun Gro Horticulture) and grown for the indicated time for each experiment. For transient expression experiments, *Nicotiana benthamiana* was grown under the same growth room conditions as *A. thaliana*, but potted in Pro-Mix PGX Biofungicide plug and germination mix.

### Quantifying powdery mildew sporulation

*G. cichoracearum* strain UCSC1 was maintained on hyper-susceptible Arabidopsis *pad4-1* mutant plants. Inoculation was carried out as described in (Serrano et al. 2014). Briefly, four-week-old plants were inoculated using a settling tower approximately 0.8 m tall and covered with a 100 micron Nitex mesh screen. Plants with a heavy powdery mildew infection (leaves covered in white powder due to production of asexual spores) were passed over the mesh to transfer the conidiospores to the plants below. Twelve *pad4-2* mutant plants were used for inoculating each tray of 60 plants. Conidiospores were counted as described in (Serrano et al. 2014). Briefly, after inoculation, the conidiospores were allowed to settle for 30 min and three leaves per genotype were harvested, weighed, and transferred to 1.5-ml microcentrifuge tubes. 500 μl of dH_2_O were added and conidiospores were liberated by vortexing 30 s at maximum speed. Leaves were removed and conidiospores were concentrated by centrifugation at 4000 g for 5 min. For each sample, conidiospores were counted in eight 1 mm^2^ fields of a Neubauer-improved haemocytometer (Marienfeld, Lauda-Königshofen, Germany). Spore counts were normalized to the initial weight of the leaves and results were averaged. The same procedure was repeated 8 days post inoculation (dpi).

### Quantifying cell death

Staining with trypan blue was performed essentially as described by (Serrano et al. 2010). Arabidopsis plants were inoculated with *G. cichoracearum* as described above, leaves collected at 5 dpi, and boiled in alcoholic lactophenol (ethanol:lactophenol 1:1 v/v) containing 0.1 mg ml^−1^ trypan blue (Sigma) for 1 min. Leaves were then destained using a chloral hydrate solution (2.5 mg ml^−1^) at room temperature overnight. Samples were observed under a Zeiss Axioplan microscope. To quantify cell death, 6 pictures of each of five experimental repetitions were randomly selected (n=30) and total leaf area and trypan-stained area were measured using ImageJ (Bethesda, MD, USA), and the percentage (area of cell death/ total leaf area) was calculated. Cell death measurements are provided as means with 10th and 90th percentiles (box) and range (whiskers).

### Plasmid construction and generation of transgenic Arabidopsis plants

EDS1^LLIF^ and PAD4^MLF^ clones used in this study were derived from pENTR cDNA clones (Bhandari et al. 2019). Site-directed mutagenesis was utilized to introduce the PAD4^S135F^ mutation into PAD4^MLF^, generating PAD4^MLFS^. All primers used in this study for cloning and site-directed mutagenesis are listed in Supplementary Table S1.

For yeast-two hybrid assays, the full-length open reading frames of EDR1, EDR1 (D810A), and EDS1 were cloned into the DNA-binding domain vector pGBKT7 (Clontech Matchmaker System). The full-length open reading frame of PAD4, PAD4^MLF^, PAD4^MLFS^, and EDS1 were cloned into the activation domain vector pGADT7. The SV40 Large T Antigen (T) and Lamin (LAM) cloned into pGADT7 and pGBKT7 respecitvely, were used as negative controls.

For yeast three-hybrid assays, EDS1 and RIN4B cDNA sequences were inserted into multiple cloning site I of the pBridge vector (Clontech) using the SmaI and SalI restriction sites (separate constructs). The EDR1 kinase domain (amino acids 587-933) and EDR1 kinase domain substrate trap mutant form (EDR1^D810A^) were cloned into multiple cloning site II of the pBridge vector using NotI and BglII restriction sites. PAD4 cDNA was inserted into the pGADT7 (Clontech) plasmid using NdeI and SmaI restriction sites. To clone AvrB into pGADT7, NdeI and BamHI restriction sites were used.

For EDR1 yeast two-hybrid experiments, EDR1 full-length wild-type cDNA and EDR1^ST^ (D810A) was cloned into pGBKT7 using SmaI and SalI restriction sites. EDS1 and PAD4 were cloned into pGADT7 using NdeI and SmaI restriction sites.

For transient expression in *N. benthamiana*, PAD4-mCherry, PAD4^S135F^-mCherry, EDS1-3xHA, EDS1^LLIF^-3xHA, and 3xHA-mCherry were cloned into the cauliflower mosaic virus 35S promoter vector pEarleyGate100 (Earley et al. 2006) using a modified multisite Gateway recombination cloning system (Invitrogen) as described in (Qi et al. 2012). PAD4-cYFP and EDS1-nYFP were cloned into the dexamethasone-inducible vectors pTA7001-GW (Aoyama and Chua 1997) and pBAV154 (Vinatzer et al. 2006), respectively, using multisite Gateway cloning. EDR1-sYFP and EDR1^ST^-sYFP were also cloned into pBAV154 using multisite Gateway cloning.

Transgenic *pad4-1* plants expressing PAD4-5xMYC, PAD4^S135D^-5xMYC, and PAD4^S135F^-5xMYC were generated using the floral dip method (Clough and Bent 1998). PAD4^S135D^ clones were generated using site-directed mutagenesis of PAD4 cDNA. PAD4, PAD4^S135D^, and PAD4^S135F^ full-length cDNA tagged with 5xMYC were cloned into the pEarleyGate100 vector (Earley et al. 2006) using multisite Gateway cloning. Plasmids were transformed into Agrobacterium strain GV3101 (pMP90) by electroporation with selection on Luria-Bertani plates containing 50 μg/mL kanamycyin sulfate (Sigma-Aldrich) and 20 μg/mL gentamycin (Gibco). Selection of transgenic plants was performed by spraying 1-week old seedlings with 300 μM BASTA (Finale). seedlings with 300 μM BASTA (Finale). Protein expression was verified via immunoblot using mouse anti-MYC-HRP antibody (ThermoFisher).

### Yeast two-hybrid and yeast three-hybrid assays

For yeast two-hybrid assays between EDR1 and PAD4 or EDS1, pGBKT7 and pGADT7 clones were transformed into haploid yeast strain AH109 (Clontech) by electroporation, and selected on SD-Trp-Leu medium. For yeast two-hybrid assays between EDS1 and PAD4, the full-length EDS1 open reading frame was cloned into an empty pBridge vector. Full-length PAD4, PAD4^S135F^, PAD4^MLF^, and PAD4^MLFS^ open reading frames were cloned into pGADT7. Yeast strain AH109 was transformed with pGADT7 vectors by electroporation and transformants were selected on SD-Leu. Yeast strain Y187 was transformed with pBridge plasmids by electroporation and transformants were elected on SD-Trp.

For yeast three-hybrid assays, EDR1-KD and EDR1-KD^ST^ were cloned into pBridge vectors, under the control of the MET25 promoter. EDS1 and RIN4B were cloned into pBridge. PAD4, PAD4^S135F^, and AvrB were cloned into pGADT7. Yeast strains AH109 and Y187 were transformed with pGADT7 and pBridge, respectively.

Matings between the Y187 and AH109 strains carrying the appropriate constructs were performed in yeast peptone dextrose medium at 30°C for 16 hours. Mating cultures were then diluted and plated on SD-Trp-Leu. Before carrying out yeast two-hybrid or three-hybrid assays, yeast were grown for 16 hours at 30°C. Cultures were re-suspended in water to an OD_600_ of 1.0, serially diluted, and plated on appropriate SD media. Plates were allowed to grow for up to 5 days at 30°C.

### β-galactosidase assays

β-galactosidase assays using ortho-Nitrophenyl-β-galactoside (ONPG) were performed as described in the Clontech Yeast Protocols Handbook 2009. Diploid yeast was grown overnight in SD–Leu-Trp at 30°C. A subculture was made by adding 4 mL of fresh SD–Leu-Trp to 1 mL of the overnight culture. The subculture was grown at 30°C until OD_600_ reached 0.3. Cells were pelleted and re-suspended in Z buffer. A 100 μL fraction was then subjected to three cycles of freezing in liquid nitrogen and thawing in a 37°C water bath. 700 μL of Z buffer containing β-mercaptoethanol was then added. 170 μL Z buffer with ONPG was then added to each reaction. Samples were incubated at 30°C for up to 24 hours. OD_600_ and OD_420_ readings were taken and β-Gal units calculated.

### Immunoprecipitations and immunoblots

For total protein extraction, four leaves of infiltrated *N. benthamiana* were collected, frozen with liquid nitrogen, and ground in lysis buffer (50 mM Tris-HCl, pH 7.5, 150 mM NaCl, 1% Nonidet P-40, 1% Plant Proteinase Inhibitor Cocktail [Sigma], and 50 mM 2,2′-Dithiodipyridine [Sigma]) or, for co-IPs, IP Buffer (50 mM Tris, pH 7.5, 150 mM NaCl, 1 mM dithiothreitol, 1mM EDTA, 1% Nonidet P-40, 10% glycerol, 1% Plant Proteinase Inhibitor Cocktail [Sigma], and 50 mM 2,2′-Dithiodipyridine [Sigma]). Samples were centrifuged at 10,000 g at 4°C for 5 minutes, and supernatants were transferred to new tubes.

Immunoprecipitations were performed as described previously (Shao et al. 2003) using GFP-Trap_A and RFP-Trap beads (Chromotek). Total proteins were mixed with 1 volume of 2× Laemmli sample buffer [Bio-Rad], supplemented with 5% β-mercaptoethanol, 1% Protease Inhibitor Cocktail [Sigma], and 50 mM 2,2′-Dithiodipyridine [Sigma]). Samples were then boiled for 5 min before loading. Total proteins and/or immunocomplexes were separated by electrophoresis on a 4-20% Mini-PROTEAN TGX Stain-Free protein gel (Bio-Rad). Proteins were transferred to a nitrocellulose membrane and probed with anti-HA-HRP (Sigma,), anti-mCherry-HRP (Santa Cruz), mouse anti-GFP (Invitrogen), and goat anti-mouse-HRP antibodies (Invitrogen).

For protein extraction from yeast, yeast grown on solid -Leu, -Trp plates were resuspended in lysis buffer (100 mM NaCl, 50 mM Tris-Cl, pH 7.5, 50 mM NaF, 50 mM Na-β-glycerophosphate, pH 7.4, 2 mM EGTA, 2 mM EDTA, 0.1% Triton X-100, 1 mM Na_3_VO_4_). Glass beads were then added to the suspension and the solution was vortexed three times for 1 minute. Samples were then boiled for 10 minutes. Immunoblots were performed using anti-HA-HRP (Sigma), mouse anti-GAL4DBD (RK5C1) (Santa Cruz Biotechnology), rabbit anti-EDS1 (Agisera), goat anti-mouse-HRP (abcam), and goat anti-rabbit-HRP (abcam) antibodies. Visualization of immunoblots from yeast strains used in three-hybrid assay were performed using the KwikQuant Imager (Kindle Biosciences).

### Transcriptome analysis

The *edr1* dataset was based upon previously generated microarray data of *edr1* plants 18 hours post inoculation with powdery mildew (Christiansen et al. 2011; GEO Accession GSE26679). Upregulated genes were identified as having higher expression in *edr1* plants compared to wildtype plants (p value < 0.05) using the NCBI GEO2R tool (Edgar et al. 2002). GENE IDs were converted to TAIR using the DAVID Gene ID Conversion Tool (Huang et al. 2008). The EDS1-PAD4 dataset was based upon 155 genes previously identified as being significantly upregulated due to EDS1 and PAD4 coexpression (Cui et al. 2017). Comparison of the *edr1* and PAD4-EDS1 datasets was performed using the Venny 2.1 tool (Oliveros 2007). Gene Ontology enrichment analysis was performed using PANTHER gene list analysis (Mi et al. 2019).

### Co-expression of EDR1, PAD4, and EDS1 for mass spectrometry

To detect phosphorylation of PAD4 or EDS1 via EDR1, PAD4-mCherry and EDS1-3xHA, were transiently co-expressed with either EDR1 or EDR1-ST(D810A)-sYFP in *N. benthamiana*. 24 hours after agrobacterium infiltration, plants were sprayed with dexamethasone to induce EDR1 and EDR1-ST expression. Immunoprecipitation and gel electrophoresis was carried out as noted above using RFP-trap (Chromotek) beads. Following gel electrophoresis, PAD4-mCherry and EDS1-HA bands were visualized using UV light, and excised. EDS1-HA and PAD4-mCherry bands were then sent for MS analysis.

Gel bands were diced into 1 mm cubes and incubated for 45 min at 57 °C with 2.1 mM dithiothreitol to reduce cysteine residue side chains. These side chains were then alkylated with 4.2 mM iodoacetamide for 1 h in the dark at 21 °C. Proteins were digested with either trypsin, chymotrypsin, or pepsin. For the trypsin digestion, a solution containing 1 μg trypsin, in 25 mM ammonium bicarbonate was added and the samples were digested for 12 hours at 37 °C. For the chymotrypsin digestion, a solution containing 1 μg chymotrypsin, in 25 mM ammonium bicarbonate was added and the samples were digested for 12 hours at 25 °C. For the pepsin digestion, a solution containing 0.5 μg of pepsin in 5% formic acid was added and the samples were digested for 12 hours at 21°C. The resulting peptides were desalted using a ZipTip (Millipore, Billerica MA). The samples were dried down and injected into an EasyNano HPLC coupled to an Orbitrap Fusion Lumos mass spectrometer (Thermo Fisher Scientific, Waltham MA) operating in data dependent MS/MS selection mode. The peptides were separated using a 75 micron, 25 cm column packed with C18 resin (Thermo Fisher Scientific, Waltham MA) at a flow rate of 300 nl/min. A one hour gradient was run from Buffer A (0.1% formic acid) to 60% Buffer B (100% acetonitrile, 0.1% formic acid).

## ACKNOWLEDGMENTS

We thank the Indiana University Light Microscopy Imaging Center for access to the Leica SP5 confocal microscope, as well as Jonathan Trinidad and Yixiang (Alex) Zhang from the Indiana University Laboratory for Biological Mass Spectrometry for performing proteomic analyses. M.N. was supported by a Carlos O. Miller Fellowship from the Indiana University Foundation. This work was funded in part by the United States National Institute of General Medical Sciences of the National Institutes of Health (Grant R01 GM063761 to R.W.I.) and by the U.S. National Science Foundation (Grant IOS-1645745 to R.W.I.). J.E.P, D.B.B and J.B. were funded by The Max-Planck Society and a Deutsche Forschungsgemeinschaft CRC 670 grant.

---

**Supplementary Fig. S1.**
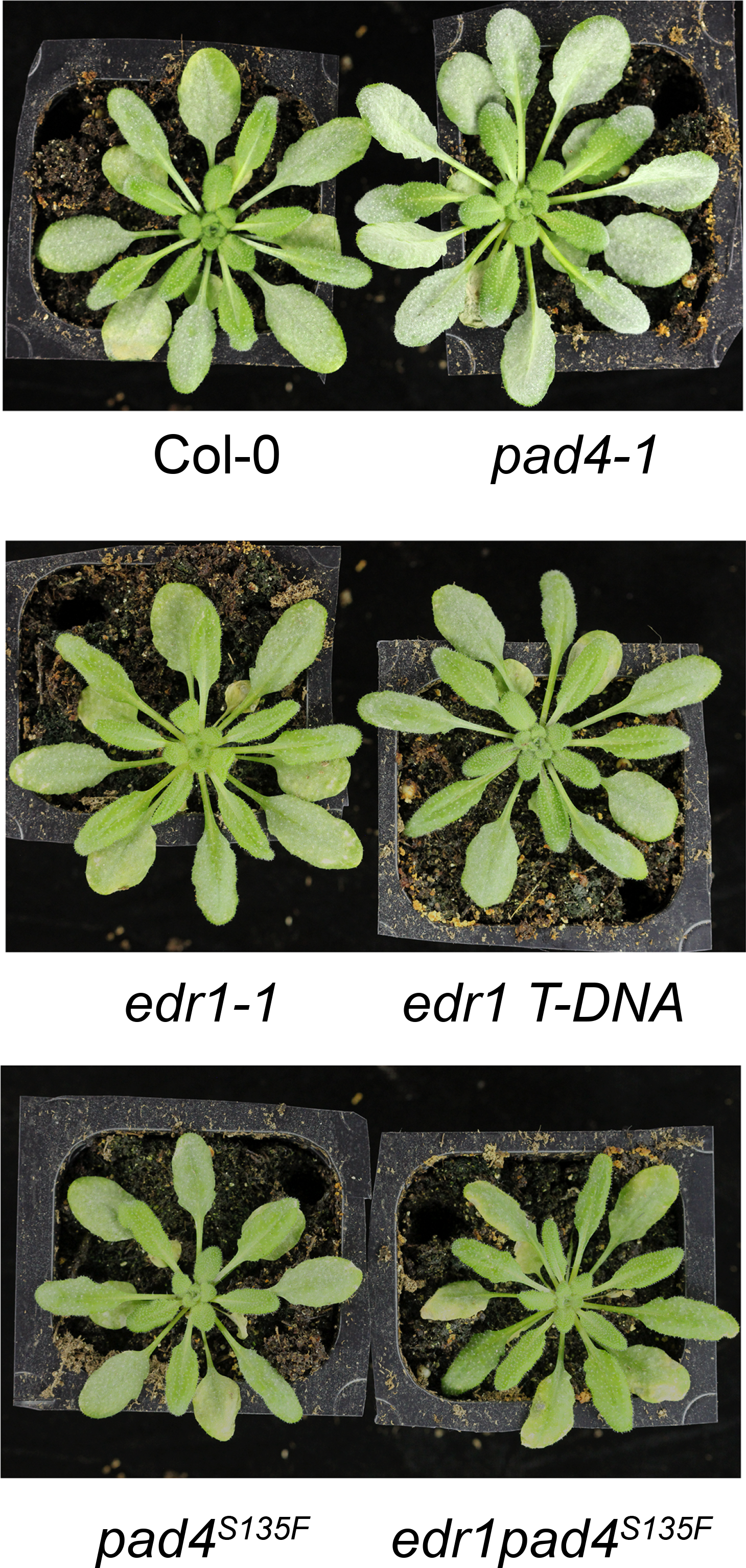
The *pad4*^*S135F*^ mutation does not result in a loss of function. **A,** Photographs of powdery mildew-infected plants 8 dpi. *pad4-1* plants display enhanced susceptibility and an increased level of powdery mildew growth, while *pad4*^*S135F*^ plants do not.

**Supplementary Fig. S2.**
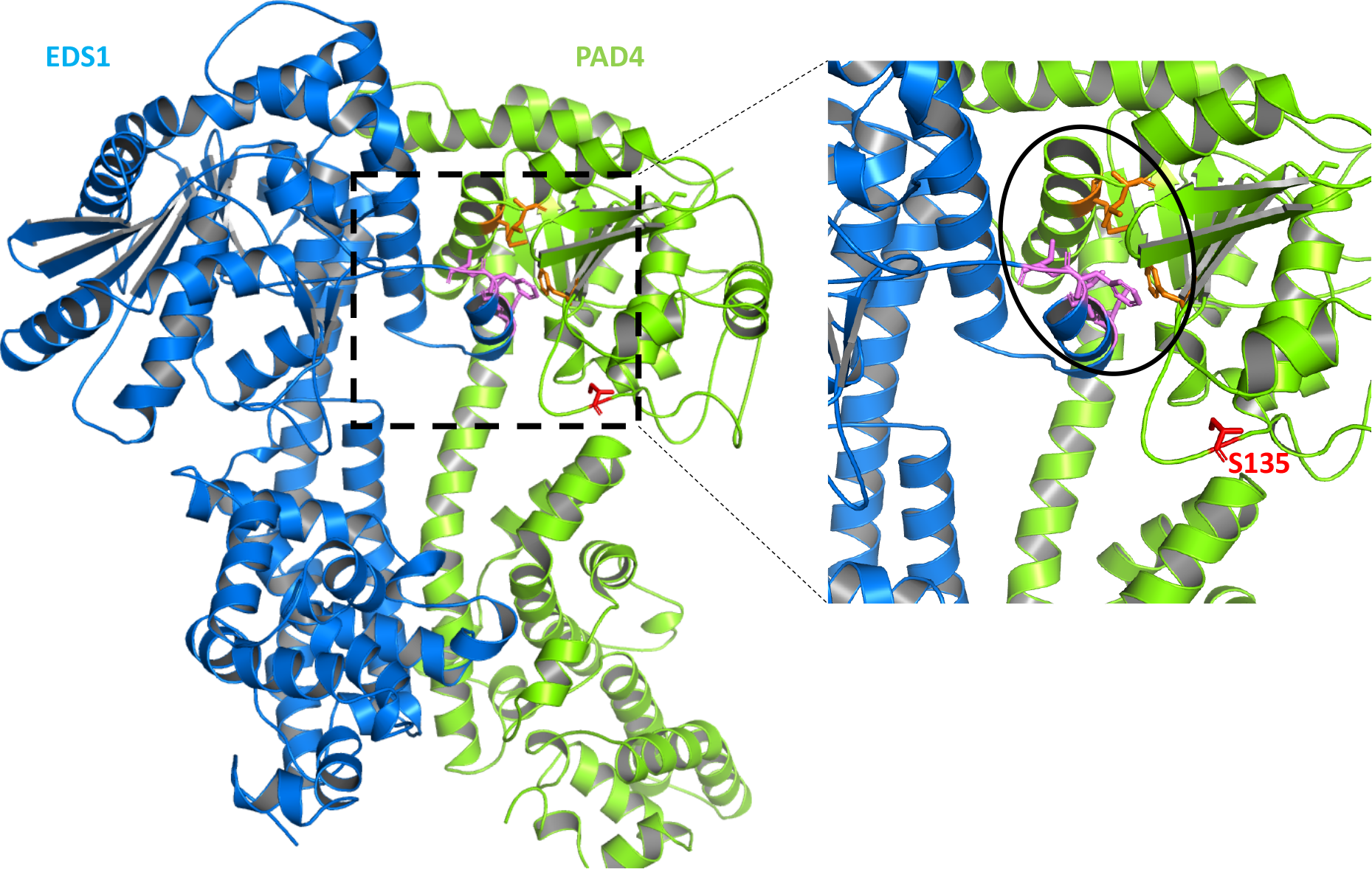
The S135F mutation in PAD4 is positioned away from the PAD4-EDS1 interaction surface. **A,** Cartoon representation of EDS1 (blue) and PAD4 (green) based on the EDS1-SAG101 structure (Wagner et al., 2013). **B,** Close-up of EDS1^LLIF^-PAD4^MLF^ hydrophobic groove mediating N-terminal binding in the heterodimer. Key N-terminal domain residues that drive heterodimerization between EDS1 and PAD4 are shown as magenta and orange sticks, respectively. PAD4^S135^ (S135, red stick) is not in direct contact with the above residues and faces away from the binding groove. The S135F mutation is therefore unlikely to interfere with EDS1-PAD4 heterodimer formation. Substitution of the PAD4 polar serine (S) residue with a bulky phenylalanine (F) at this position might, however, cause structural reorganization that could affect EDS1-PAD4 signaling.

**Supplementary Table S1.**
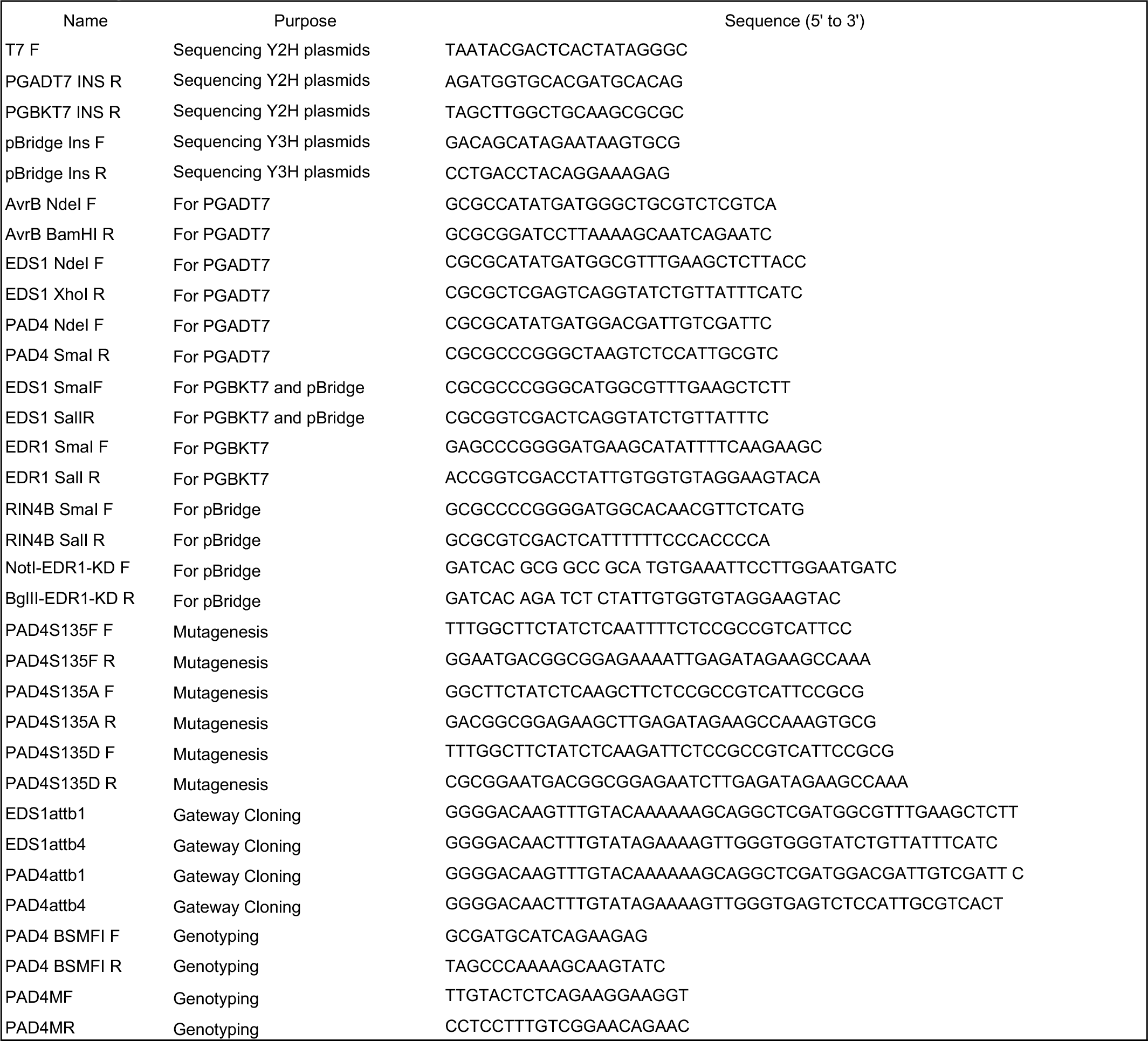
Primers used in this study.

